# VERICIGUAT RESCUES CYCLIC GUANOSINE MONOPHOSPHATE PRODUCTION IN HUMAN AORTIC VASCULAR SMOOTH MUSCLE CELLS AND AUGMENTS VASORELAXATION IN AORTIC RINGS EXPOSED TO HIGH GLUCOSE

**DOI:** 10.1101/2024.06.21.600154

**Authors:** David Polhemus, Diego Almodiel, Tarek Harb, Efthymios Ziogos, Nuria Amat-Codina, Mark Ranek, Lakshmi Santhanam, Gary Gerstenblith, Thorsten Leucker

**Affiliations:** Division of Cardiology, Department of Medicine, Johns Hopkins University, Baltimore, MD; Department of Chemical & Biomolecular Engineering, Johns Hopkins University, Baltimore, MD; Department of Anesthesiology & Critical Care Medicine, Johns Hopkins University, Baltimore, MD

**Keywords:** Nitric oxide, cGMP, inflammation, vascular smooth muscle cell function, protein kinase G activity

## Abstract

**Background:** Normal endothelial cell dependent vascular smooth muscle cell function is mediated by nitric oxide (NO), which stimulates soluble guanylyl cyclase (sGC) production of the second messenger, cyclic guanosine monophosphate (cGMP) leading to increased protein kinase G (PKG) activity and vascular smooth muscle relaxation. NO bioavailability is impaired in inflammatory settings, such as high glucose (HG). We examined whether the direct sGC sensitizer/stimulator vericiguat, augments cGMP production in human vascular smooth muscle cells (HVSMC) exposed to high glucose and explored its effect on vasorelaxation.

**Methods:** Aortic HVSMCs were exposed to HG for 24h. In the treatment group, cells also received 1uM vericiguat for 24h. After incubation, cGMP and PKG activity were measured. Additionally, thoracic murine aortas were exposed to HG or to normal glucose (NG) control. The rings were then placed in an organ chamber bath and dose response curves to increasing doses of acetylcholine (Ach) and sodium nitroprusside were constructed for three groups: control (normal glucose), HG alone, and HG + vericiguat.

**Results:** HVSMCs exposed to HG produced significantly less cGMP than those exposed to NG. cGMP production in the presence of HG was rescued when treated with 1uM vericiguat. Additionally, PKG activity was impaired in the presence of HG and enzyme activity was restored with vericiguat. In isolated mouse aortic rings, ACh mediated relaxation was impaired following treatment with HG, but was improved when a HG group was treated with vericiguat.

**Conclusions:** The sGC sensitizer/stimulator vericiguat restored cGMP production and PKG activity in the setting of HG. Vericiguat enhanced ACh-mediated vasorelaxation in the setting of HG. The findings suggest clinical studies are warranted to investigate the potential of sGC sensitization/stimulation as a therapeutic intervention to improve vascular endothelial-dependent function that is impaired in pro-inflammatory settings that are associated with the development of atherosclerotic disease.

## INTRODUCTION

Blood flow regulation and vascular homeostasis are centered on optimal communication between, and function of, the endothelial cells (EC) and vascular smooth muscle cells (VSMC). A central homeostatic mediator in this system is nitric oxide (NO). NO is a gaseous signaling molecule that is generated and released by EC, diffuses into VSMC, and exerts potent vasodilatory, pro-survival, anti-inflammatory, and anti-hypertrophic effects via its ability to increase soluble guanylate cyclase (sGC) activity and ultimately the production of the second messenger cyclic guanosine monophosphate (cGMP)^1,2^. Furthermore, cGMP binding activates protein kinase G (PKG), which phosphorylates serines and threonines on substrate proteins. Endothelial cell-derived NO bioavailability is diminished, however, in pro-inflammatory states such as hyperglycemia, metabolic syndrome, and aging^3^. This in turn results in impaired VSMC cGMP signaling. cGMP has emerged as a novel cardiovascular therapeutic target, recently attracting attention for the potential management of pulmonary hypertension, heart failure, and cardiac injury post myocardial infarction^4–6^. An intervention which increases sGC activity directly, or by increasing the response to a given NO stimulus (i.e. sensitizing sGC), offers the opportunity to overcome inflammation-induced vascular dysfunction due to the decreased NO bioavailability present in most pro-inflammatory states^7,8^. A class of compounds called sCG sensitizers/stimulators bind to sGC and promote cGMP production. One such compound, vericiguat, was evaluated in the VICTORIA trial and was found to reduce the incidence of death from cardiovascular causes or hospitalization in high-risk heart failure patients^9^. Given that dysfunctional NO-cGMP signaling contributes to impaired vascular function in pro-inflammatory states, we investigated vericiguat’s ability to restore cGMP signaling and vascular function in *in vitro* and *ex vivo* models of high glucose, a pro-inflammatory setting.

## METHODS

### Isolated Human Aortic Vascular Smooth Muscle Cell Studies

Human Aortic VSMCs (Lonza Cat No CC-2571 batch no: 21LT316170) were placed in 6 well places. Once cells reached 60%-70% confluency, the media was changed to a serum free media. SmGM-2 Smooth Muscle Cell Growth (Lonza Cat No. CC 3182) was used for incubation. After 24 hours, cells were treated with three different conditions as follows. Cells were exposed to normal glucose (5.5 mM D-glucose and 24.5 mM L-glucose), high glucose (30mM D-glucose), or to high glucose + vericiguat 1uM for 24 hours. All groups were treated with sildenafil (1uM, TCI America and Cat No. S098625MG) and the NO donor, DETA NONOate (0.1 uM, Cayman Chemicals and Cat No 82120). The NO donor was given because this VSCM system was void of endothelial cells (and thus endogenous NO production). Sildenafil was added to the system to increase cGMP half-life and facilitate assay measurement of the otherwise unstable compound. After 24 hours of incubation, cells were isolated, lysed (Cell signaling Lysis Buffer #9803), and then used to measure cGMP and PKG activity.

### Vericiguat Preparation

A 2.5 mg tablet of vericiguat was dissolved in Tris buffered saline (TBS) resulting in a final concentration of 540.2 uM. The vericiguat stock solution was further diluted for the downstream *in vitro* assays.

### Cyclic Guanosine Monophosphate (cGMP measurement)

Following cell isolation and lysis, intracellular cGMP was measured by ELISA according to the manufacturer’s instruction (Cell Signaling Cat No 4360).

### Protein Kinase G activity assay

In vitro PKG1 activity was assessed by in vitro colorimetric assay (Cyclex, CY-1161, MBL International) following the manufacturer’s instructions as previously described^10 11^. The assay provides cGMP substrate and a kinase-specific peptide target to assess phosphorylation activity.

### Aortic Ring Vascular Reactivity

Male C57Bl/6J mice at the age of 8-12 weeks were purchased from Jackson Labs. All procedures involving the use of animals were approved by the Institutional Animal Care and Use Committee. Mice were anesthetized with isoflurane at a 3% concentration until unresponsive to stimuli. Thoracic aortas were carefully dissected and excised for ex vivo vascular reactivity testing (myograph technology) as previously described^12,13^. Aortic ring segments were exposed to normal glucose (11mM) or high glucose (40mM) in Krebs-Henseleit buffer and incubated for 24 hours at 4 degrees C. Rings were pre-contracted with phenylephrine and treated with escalating doses of acetylcholine (10-9 to 10-5M) or nitroprusside (10-9 to 10-5M) after receiving 10^-5.5 uM vericiguat or control. A dose response curve was constructed for escalating dosing of vericiguat (10-9 to 10-3.5M).

### Statistical Analyses

All data are presented as mean ± standard error of the mean (SEM). All statistical analyses were performed using Prism 10 software (GraphPad, San Diego, CA). For multiple group comparisons, 1-way ANOVA was used. For data sets involving multiple factors, such as multiple concentrations (e.g. vascular reactivity experiments), 2-way ANOVA followed by multiple comparison testing was used. A p-valve of <0.05 was considered statistically significant.

## RESULTS

### Vericiguat rescues cGMP production and signaling in the setting of high glucose

Figure 1A provides an overview of the proposed mechanisms. Various doses of vericiguat were examined in isolated human aortic vascular smooth muscle cells. A dose of 1uM was utilized for the following experiments because it was the lowest tested concentration that led to a measurable increase in cGMP in our system (Figure 1B). VSMCs were incubated with high glucose (30mM), which led to impaired cGMP production (Figure 1C). However, in the setting of high glucose, vericiguat restored cGMP production to that of the control group with normal glucose levels. PKG activity was impaired in VSMCs exposed to high glucose, but enzyme activity was restored in the high glucose cells treated with vericiguat (Figure 1D).

**Figure 1:**
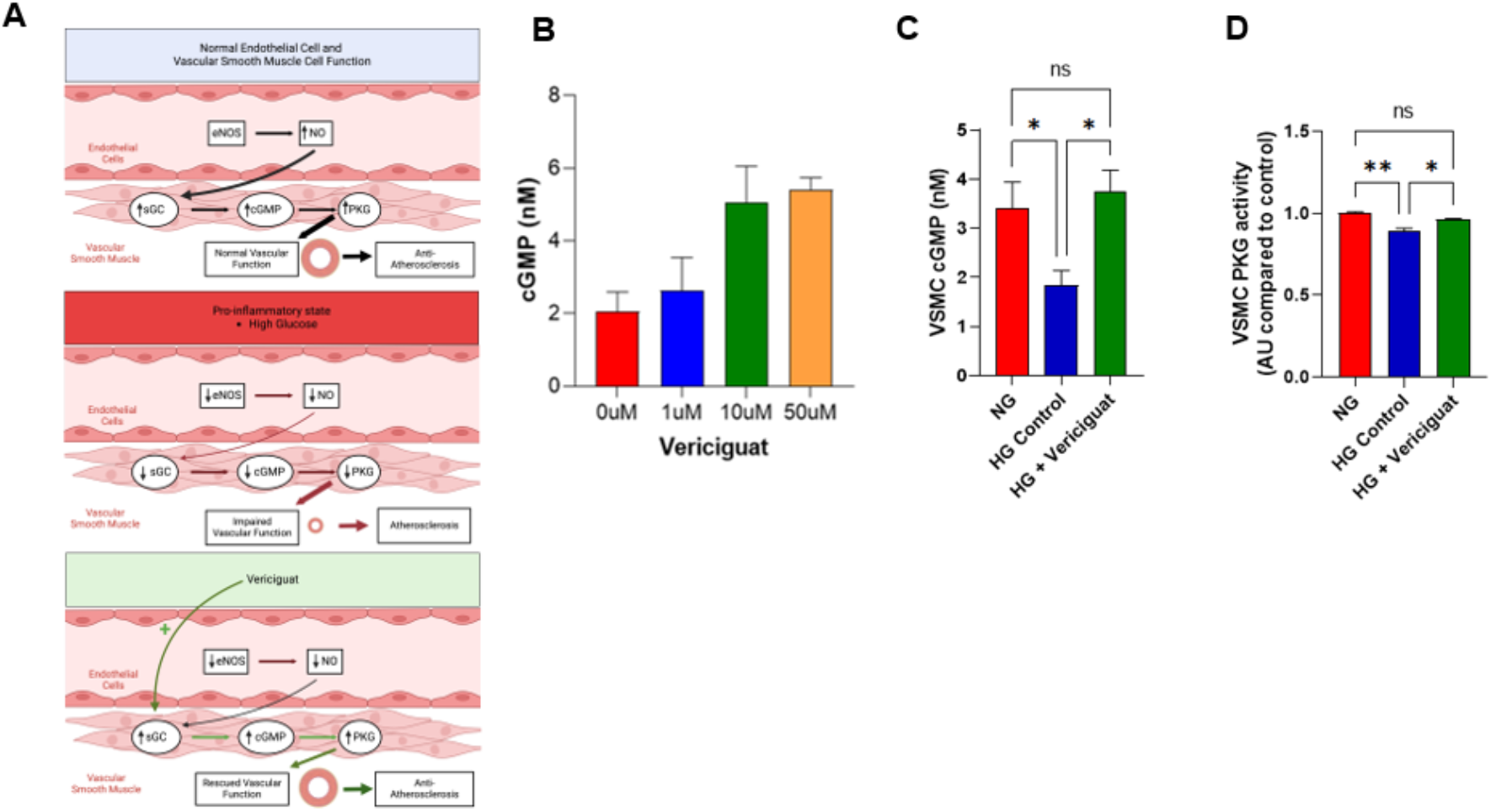
cGMP signaling is restored by vericiguat in human aortic VSMCs exposed to high glucose. (A) Proposed mechanism of eNOS-NO-cGMP signaling in healthy and pro-inflammatory states and the actions of vericiguat to bypass NO production and restore cGMP-PKG signaling in NO deficient states (B) cGMP production by human aortic VSMCS at various doses of vericiguat in a normal glucose concentration system. (C) cGMP production of VSMCs subjected to normal glucose, high glucose, and high glucose +1µM Vericiguat. *p<0.05 (n=9 per group) by 1 way ANOVA. (D) Protein Kinase G (PKG) activity of VSMCs subjected to normal glucose, high glucose, and high glucose +1µM Vericiguat. *p<0.05, **p<0.01 by 1 way ANOVA. (n=3 per group)

### Vericiguat improves endothelial dependent vascular reactivity in a high glucose setting

Vascular reactivity was assessed in aortic rings in the presence of normal glucose of high glucose. First, a vericiguat dose response curve was constructed in rings exposed to normal and high glucose (Figure 2A). Vericiguat concentration of 10^-5.5M was used in the subsequent acetylcholine (Ach) and sodium nitroprusside (SNP) studies as it caused only ∼5% maximal relaxation. Endothelial-dependent Ach mediated relaxation was impaired in aortic rings expose to high glucose (Figure 2B). Interestingly, treatment with vericiguat prior to escalating doses of Ach in aortic rings exposed to high glucose not only augmented vascular reactivity compared to the high glucose group, but augmented vasorelaxation compared to the normal glucose group (Figure 2B). Endothelial-independent SNP mediated vasorelaxation was not impaired in aortic rings exposed to high glucose and pre-treatment with vericiguat did not enhance SNP mediated vasorelaxation (Figure 2D-E).

**Figure 2:**
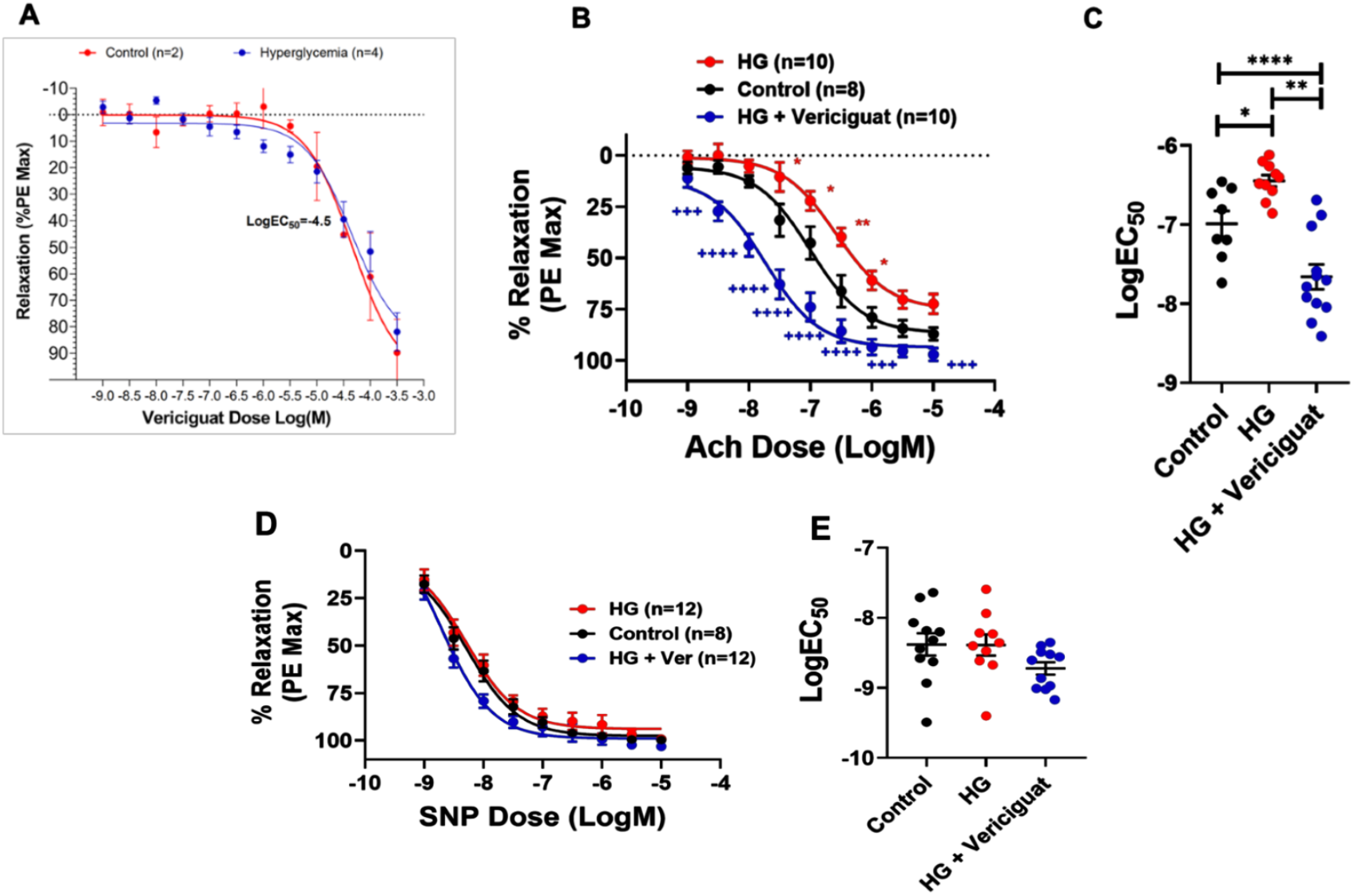
Vericiguat restores endothelial mediated vasorelaxation in the setting of high glucose. (A) Dose response curves of vericiguat in aorta pre-treated with high glucose or normal glucose. (B) Ach dose response curve of vessels subjected to control, high glucose, and high glucose + vericiguat *p<0.05 vs control by one-way ANOVA. **p<0.01 vs control by one-way ANOVA. +++p<0.001 vs HG by one-way ANOVA. ++++ p<0.001 vs HG by two-way ANOVA. (C) LogEC50 of Ach dose response curves in figure B. *p<0.05, **p<0.01, ****p<0.0001 by one-way ANOVA. (D) SNP dose response curves of aorta subjected to control, high glucose, and high glucose + vericiguat. (E) LogEC50 of SNP dose response curves.

## DISCUSSION

Vascular endothelial cell dysfunction is a defining characteristic of much of vascular pathobiology and is often associated with redox imbalance, endothelial nitric oxide synthase (eNOS) dysfunction, and impaired NO signaling^14^. Pro-inflammatory states and environments of high oxidative stress impair eNOS-NO signaling. For example, a decrease in NO bioavailability caused by hypercholesterolemia-induced endothelial dysfunction was shown to impair vasodilation^15,16^. Additionally, NO inhibits endothelial cell activation and adhesion that occur in response to inflammatory mediators^17^. NO also has been shown to have anti-atherosclerotic effects, anti-inflammatory actions, and plays a role in maintaining endothelial integrity^18,19^ and models of late stage atherosclerosis are associated with impaired cGMP-PKG signaling^20^.

The signaling pathway of NO is complex, with both paracrine and autocrine functions. It plays an important regulatory role in vascular homeostasis and has been a target for pharmacotherapy since its discovery as a key mediator in vasodilation^21^. Designing NO based therapies that can continuously deliver NO pharmacologically at therapeutic levels has several challenges. The half-life of NO is incredibly short given its chemical instability. Due to its gaseous properties and its ability to freely traverse cell membranes, it can also react indiscriminately with a narrow therapeutic index before leading to the generation of reactive oxygen species and peroxynitrite^22^. The development of NO donors, which serve as pharmacologically active carriers of NO, are designed with the intention of delivering NO to target tissues. Such compounds have been designed to release NO by environmental factors such as pH, local enzyme activity, light and temperature^23^. However, the use of many NO donors designed to serve as pharmacologically active carriers of NO, such as SNP, is limited to the acute setting as their actions are short lived. The use of long-acting nitrates is limited by the development of tachyphylaxis whereas calcium channel blockers, although they may decrease inappropriate coronary vasoconstriction, lack the anti-inflammatory, anti-proliferative, and other potential anti-atherosclerotic benefits of NO.

In light of these limitations, a newer class of compounds, classified as novel oral soluble guanylate cyclase sensitizers/stimulators were created^24^. They bypass eNOS-NO generation by binding directly to a NO-independent binding site on sGC to facilitate continuous production of cGMP. One such compound, vericiguat, was clinically tested for the treatment of chronic heart failure as reported in the VICTORIA trial in 2020^9^. Treatment with vericiguat led to a significant reduction in the primary-outcome event rate of composite death from cardiovascular cause or heart failure hospitalization (hazard ratio 0.9 and p=0.02).

The aim of this study was to examine, in the preclinical setting, whether an sGC sensitizer/stimulator could restore vascular smooth muscle cGMP production and endothelial-dependent VSMC relaxation that was impaired in the setting of high glucose, an inflammatory stimulus associated with decreased NO bioavailability. Specifically, we examined vericiguat in pro-inflammatory *in vitro* and *ex vivo* vascular smooth muscle cell systems. As reported above, vericiguat effectively restored impaired cGMP signaling in human aortic VSMCs exposed to a pro-inflammatory insult. The non-endothelial dependent SNP relaxation response curve was not altered with high glucose, indicating that unlike endothelial-dependent VSMC function, VSMC function is not impaired by high glucose. The clinical implications of these findings extend to the targeting of those vascular disease states associated with inflammation. Specifically, these findings have potential application for the treatment of atherosclerotic disease whose pathology and progression are linked to heightened inflammation and impaired NO-cGMP signaling^25–27^. Inflammation has been shown to mediate all stages of the development of atherosclerosis from initiation to the associated atherothrombotic complications. Endothelial dysfunction is an early event in atherogenesis and constitutive endothelial NO release preventing leukocyte adhesion and smooth muscle proliferation, important stages in the development of atherosclerosis^27^.

In summary, we report that the sGC sensitizer/stimulator vericiguat restores cGMP-PKG signaling and vascular function in pro-inflammatory human *in vitro* and murine *ex vivo* systems. Future studies are needed to test whether these findings extend to animal and human models of inflammation-associated vascular diseases, such as diabetes.

## DISCLOSURES

This work was funded by Merck. Merck, the funder of the study, had no role in the design of the study, the collection, management, or interpretation of the data, or the statistical analysis. The funder reviewed the first submitted version of the article but was not involved in the writing or approval of the article or the decision to submit the article for publication. This work was also supported by the NIH with R01HL148112 01 to Dr. Lakshmi Santhanam

## Notes

### Competing Interest Statement

The authors have declared no competing interest.

